# Structure of human CST-pol-α/primase bound to a telomeric overhang poised for initiation of telomere C-strand synthesis

**DOI:** 10.1101/2021.12.16.472968

**Authors:** Qixiang He, Xiuhua Lin, Bianca L. Chavez, Benjamin L. Lusk, Ci Ji Lim

## Abstract

Telomere replication and regulation protect mammalian chromosome ends and promote genome stability. An essential step in telomere maintenance is the C-strand fill-in process, which is the *de novo* synthesis of the complementary strand of the telomere overhang. This step is catalyzed by polymerase-alpha/primase complex (pol-α/primase) and coordinated by an accessory factor, CTC1-STN1-TEN1 (CST). Using cryogenic-electron microscopy single-particle analysis, we report the structure of the human telomere C-strand fill-in preinitiation complex (PIC) at 3.9 Å resolution. The structure reveals a CST and a pol-α/primase co-bound to a single telomere overhang, poised for *de novo* RNA primer synthesis. Upon PIC assembly, the pol-α/primase undergoes large conformation change from its apo-state; CST partitions the DNA and RNA catalytic centers of pol-α/primase into two separate domains and positions the 3*′* end of an extended telomere single-stranded DNA template towards the RNA catalytic center (PRIM1 or p49). The telomeric single-stranded DNA template is further positioned by the POLA1 (or p180) catalytically dead exonuclease domain. Together with CST, the exonuclease domain forms a tight-fit molecular tunnel for template direction. Given the structural homology of CST to Replication Protein A (RPA), our structure provides the structural basis for a new model of how pol-α/primase lagging-strand DNA synthesis is coordinated by single-stranded DNA-binding accessory factors.

## Main

The DNA polymerase-alpha/primase complex (pol-α/primase) is needed to initiate DNA synthesis during genome replication^1,2^. The four-subunit pol-α/primase (PRIM1-PRIM2-POLA1-POLA2, also known as p49-p58-p180-p70) does this by using its primase domain to *de novo* synthesize a short 7-11 nt RNA primer, which is then extended by its DNA polymerase domain^1,2^. In cells, pol-α/primase requires modulation by highly-conserved single-stranded DNA-binding proteins (accessory factors) such as the replication protein A (RPA)^3^ and CTC1-STN1-TEN1 (CST)^4^ complexes. Both of these protein complexes share functions^5,6^ and structural homology^7^. At mammalian telomeres, CST inhibits extension of telomere G-rich overhangs^8,9^ and facilitates pol-α/primase to synthesize the complementary C-strand. The latter process is known as the telomere C-strand fill-in^10,11^. In addition, the CST-pol-α/primase machinery is essential for restarting stalled DNA replication forks genome wide and regulating end resectioning at double-stranded DNA breaks^8,12-14^. Despite a central role in eukaryotic DNA metabolism, structural insight into how pol-α/primase and its accessory factors perform their functions is lacking^1,2^.

Here, we report a 3.9 Å resolution cryogenic-electron microscope (cryo-EM) structure of the human CST-pol-α/primase co-complex bound to a telomeric single-stranded DNA (ssDNA). This represents a preinitiation state of the human telomere C-strand fill-in machinery. The structure reveals that pol-α/primase interacts with CST at two separate sites, segregating its RNA and DNA catalytic centers. CST binds most of the telomere ssDNA template and directs its 3’ end of the ssDNA towards the RNA catalytic center of PRIM1, which is poised for initiating *de novo* RNA synthesis. The catalytically dead exonuclease domain of POLA1 interacts with CTC1 and the STN1 C-terminal domain to form a tight-fit molecular tunnel for the ssDNA template to thread through. Collectively, the architecture of the telomere C-strand fill-in preinitiation complex reveals how CST, an accessory factor of pol-α/primase, organizes the enzyme for *de novo* RNA primer synthesis and provides a molecular model for the subsequent RNA-to-DNA synthesis switch.

## The architecture of human telomere C-strand fill-in preinitiation complex

To determine the structure of the human CST-pol-α/primase co-complex using cryo-EM single-particle analysis^15,16^, we expressed and purified recombinant human CST and pol-α/primase complexes and used these separately purified complexes to reconstitute the 7-subunit co-complex – CTC1-STN1-TEN1-PRIM1-PRIM2-POLA1-POLA2 (**Fig. 1a and Supplementary Fig. 1a**). As the intrinsically disordered POLA1_1-337_ domain interacts with other replication factors^3,17^, we tested if CST directly interacts with pol-α/primase via this domain. We found that both complexes directly interact with each other, independent of nucleic acids or POLA1_1-337_ (**Supplementary Fig. 1b**). To reconstitute the co-complex, we used an excess of CST and then enriched for the co-complex by pol-α/primase pull-down (**Fig. 1b**). Our attempts to solve the apo-state structure of the co-complex were unsuccessful. Two-dimensional (2D) classification analysis of negative-stain EM images revealed the apo-state co-complex had a heterogeneous conformation (**Supplementary Fig. 2a**). Interestingly, CST and pol-α/primase are homogeneous in their respective conformations when individually analyzed (**Supplementary Fig 2b & c**).

**Fig. 1:**
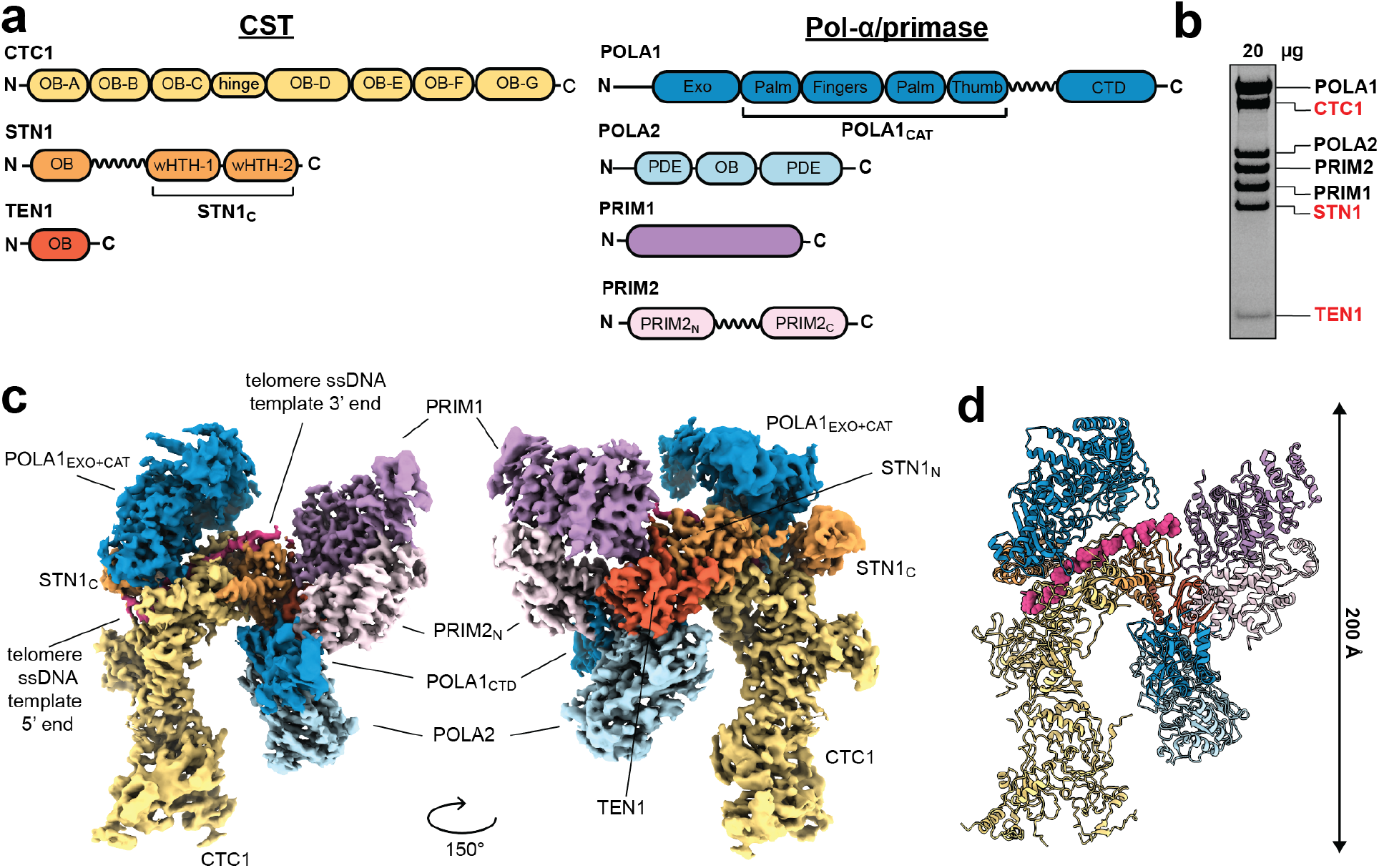
The architecture of human telomere C-strand fill-in preinitiation complex. **a**, Schematics of CTC1-STN1-TEN1 (CST) and DNA polymerase-alpha/primase complex (pol-α/primase) domains. Zigzag lines indicate flexible linkers connecting domains. **b**, Coomassie-stained denaturing protein gel of purified reconstituted CST-pol-α/primase co-complex. Black labels are pol-α/primase subunits and red labels are CST subunits. **c**, Views of the cryo-EM map of the CST-pol-α/primase preinitiation complex (PIC). **d**, Ribbon representation of the atomic model built from the cryo-EM map in **c**. For **c, d**, each domain is colored as in **a**.

To stabilize the co-complex for cryo-EM structural determination, we screened for a minimal length ssDNA of telomeric sequence that both CST and pol-α/primase can bind. Our DNA-binding gel-shift assays show pol-α/primase binds telomeric single-stranded DNA (ssDNA) less stably than CST (**Supplementary Fig. 3a-c**). As previously shown^5,8^, CST needs a minimum of three telomeric repeat to bind tightly and in the case of pol-α/primase, any detectable binding. The differential in binding affinity to telomeric ssDNA between CST and pol-α/primase are more than ten-fold for three- or four telomeric repeat ssDNA. But with a longer telomeric ssDNA (six-repeat), the gap is smaller, indicating pol-α/primase binding affinity increases with template length.

Negative-stain EM 2D classification analysis revealed that the addition of telomeric ssDNA to CST-pol-α/primase co-complex resulted in a new homogeneous conformation that was different from CST or pol-α/primase alone (**Supplementary Fig. 4**). We only saw the new conformation when telomere ssDNA of 3-repeat or longer was used, consistent with our binding assay results (**Supplementary Fig. 3**). This new conformation suggests that CST-pol-α/primase co-complex binding to telomeric ssDNA (≥ 3xTEL) improves its conformational stability. Using cryo-EM single-particle analysis, we solved the structure of the human CST-pol-α/primase co-complex bound to a telomeric ssDNA template [C3xTEL, C(TTAGGG)_3_] at a global resolution of 3.9 Å (**Fig. 1c, Supplementary Fig. 5, Supplementary Table 1**). The map resolution allows us to dock all seven subunits of CST and pol-α/primase (from AlphaFold models^18^), and the ssDNA template in the cryo-EM map, revealing the architecture of the human C-strand fill-in preinitiation complex (PIC) (**Fig. 1d**).

### Pol-α/primase separates into polymerase and primase domains upon CST-ssDNA binding

Multiple key insights are gained from the structure to understand how CST and pol-α/primase assembly on a telomeric ssDNA coordinates C-strand fill-in. The apo-state architecture of pol-α/primase changes upon assembly with CST and telomeric ssDNA, from a compact structure to a segregated form (**Fig. 2a and Supplementary Fig. 6a**). The conformational change leads to separation of the RNA^19^ and DNA^20^ catalytic centers of pol-α/primase, in the PRIM1 and POLA1 subunits, respectively (**Fig. 2b**). This also exposed the POLA1 DNA catalytic center, which was buried in the pol-α/primase apo-state structure^2,21^.

**Fig. 2:**
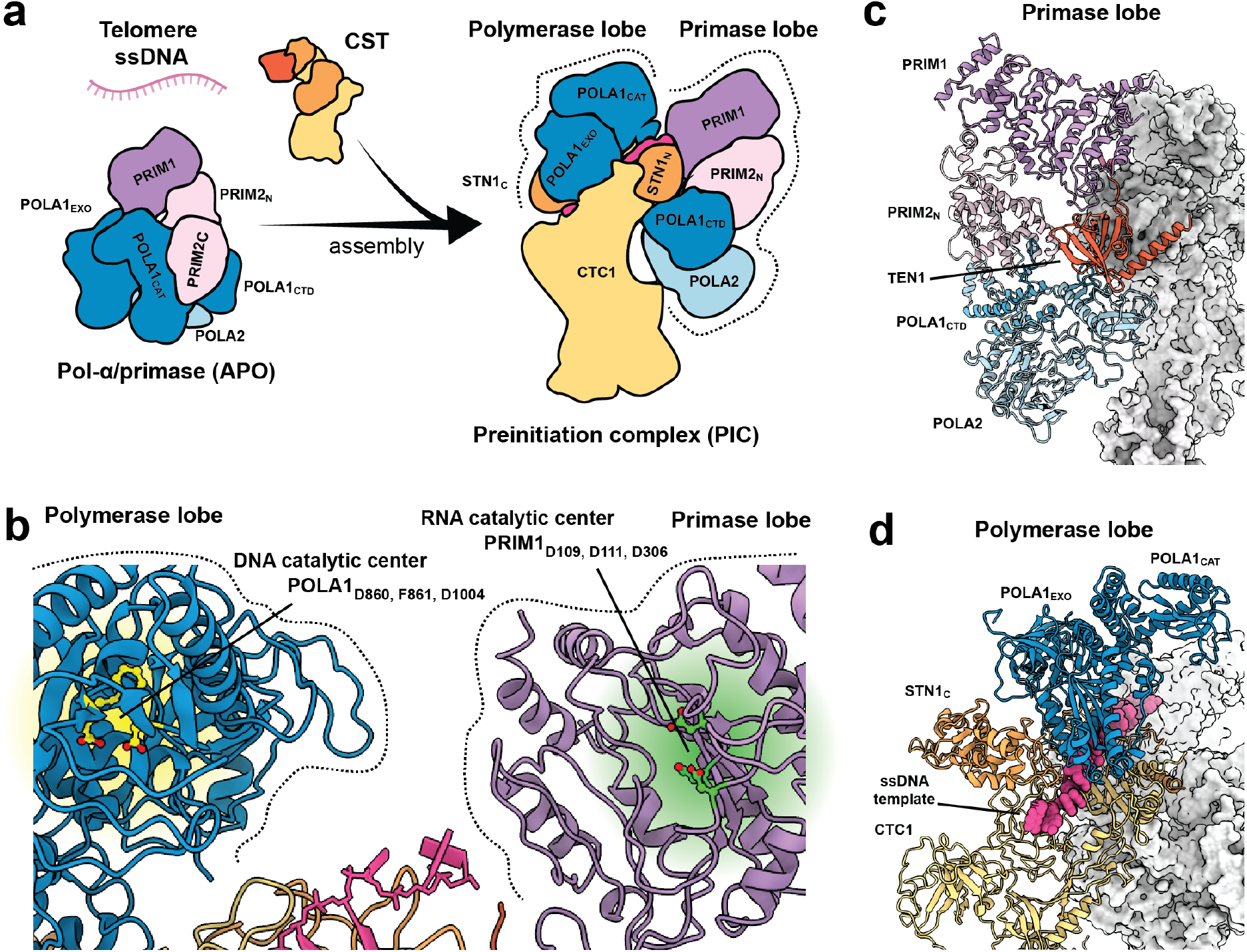
Pol-α/primase separates into polymerase and primase domains upon CST-ssDNA binding. **a**, Preinitiation complex (PIC) assembly model where pol-α/primase undergoes large conformational changes upon binding to the CST-ssDNA complex. In the PIC, pol-α/primase is spatially separated into primase and polymerase functional lobes. The lobes are encircled by dashed lines. The flexible linker connecting POLA1_CAT_ and POLA1_CTD_ domains and the PRIM2_C_ domain are not shown in this cartoon drawing. The APO pol-α/primase cartoon model is adapted from PDB 5EXR. The CST cartoon model is adapted from PDB 6W6W. **b**, Ribbon model overview of the DNA and RNA catalytic centers of POLA1 and PRIM1 respectively. Amino acid residues underlying the DNA and RNA catalytic centers are colored yellow and green respectively. The pink-colored molecule is the modeled single-stranded DNA. **c**, TEN1 subunit interactions with all four subunits (PRIM1, PRIM2_N_, POLA1_CTD_ and POLA2) of the primase lobe defined in **a**. The rest of the PIC is shown as white-colored surface representation. **d**, CTC1, ssDNA and STN1_C_ interact with POLA_EXO_ to support the polymerase domain defined in **a**. The rest of the PIC is shown as white-colored surface representation. Colors are defined similarly in all panels: CTC1 in yellow, STN1 in orange, TEN1 in dark orange, PRIM1 in purple, PRIM2 in light purple, POLA1 in blue, and POLA2 in light blue.

CST served as a scaffold to mediate this pol-α/primase conformational change. The STN1_n_-TEN1 domain of CST binds the “primase lobe” (PRIM1-PRIM2_N_-POLA1_CTD_-POLA2) of pol-α/primase using multiple contact sites; TEN1 binds all four subunits of the primase lobe and the protruding PRIM1_85-97_ loop sits in the interface between STN1_n_ and TEN1 (**Fig. 2c and Supplementary Fig. 6b**). These interactions provide the molecular basis as to why STN1-TEN1, in the context of a CST heterotrimer, is essential for telomere C-strand fill-in process^22-24^. This also defines a new structure-function role for TEN1.

The CTC1_OB-F_ and STN1_C_ domains of CST bind the POLA1_EXO_ of the “polymerase lobe” (POLA1_EXO_ and POLA1_CAT_) (**Fig. 2d**) - STN1_C_ sits between CTC1_OB-F_ and POLA1_EXO_. This new STN1_C_ docking position, in addition to the two other positions that control CST decamerization^7^, underlines STN1_C_ function as an allosteric switch to regulate CST inter-and intra-molecular interactions. The “thumb” subdomain of POLA1_CAT_ and PRIM2_C_ domain of pol-α/primase are highly flexible, as indicated by local resolution analysis of the cryo-EM map (**Supplementary Fig. 7**).

### CST positions the ssDNA template for pol-α/primase RNA primer synthesis

The cryo-EM map density of the ssDNA template in our structure corroborates the previously determined CST ssDNA-binding anchor site^7^. It also reveals how CST binds ssDNA beyond its anchor site. As the local resolution of the ssDNA map density is ∼ 3.4 Å (**Supplementary Fig. 7**), we used the previously modeled 4-nt telomeric ssDNA^7^ as a starting reference to build our 15 nt ssDNA model (guided by optimal cross-correlation value). CST binds the telomeric ssDNA template (**Fig. 3a)** and directs the template 3’ end towards the RNA catalytic center of pol-α/primase (**Fig. 3b)**. In addition to the DNA-binding residues (in CTC1_OB-F_ and flexible loop of CTC1_OB-G_) of the ssDNA-binding anchor site^7^, CST uses the β-hairpin loop of CTC1_OB-G_ (CTC1_1175-1190_) and STN1_N_ to bind the 3’ end portion of the ssDNA template (**Supplementary Fig. 8a**).

**Fig. 3:**
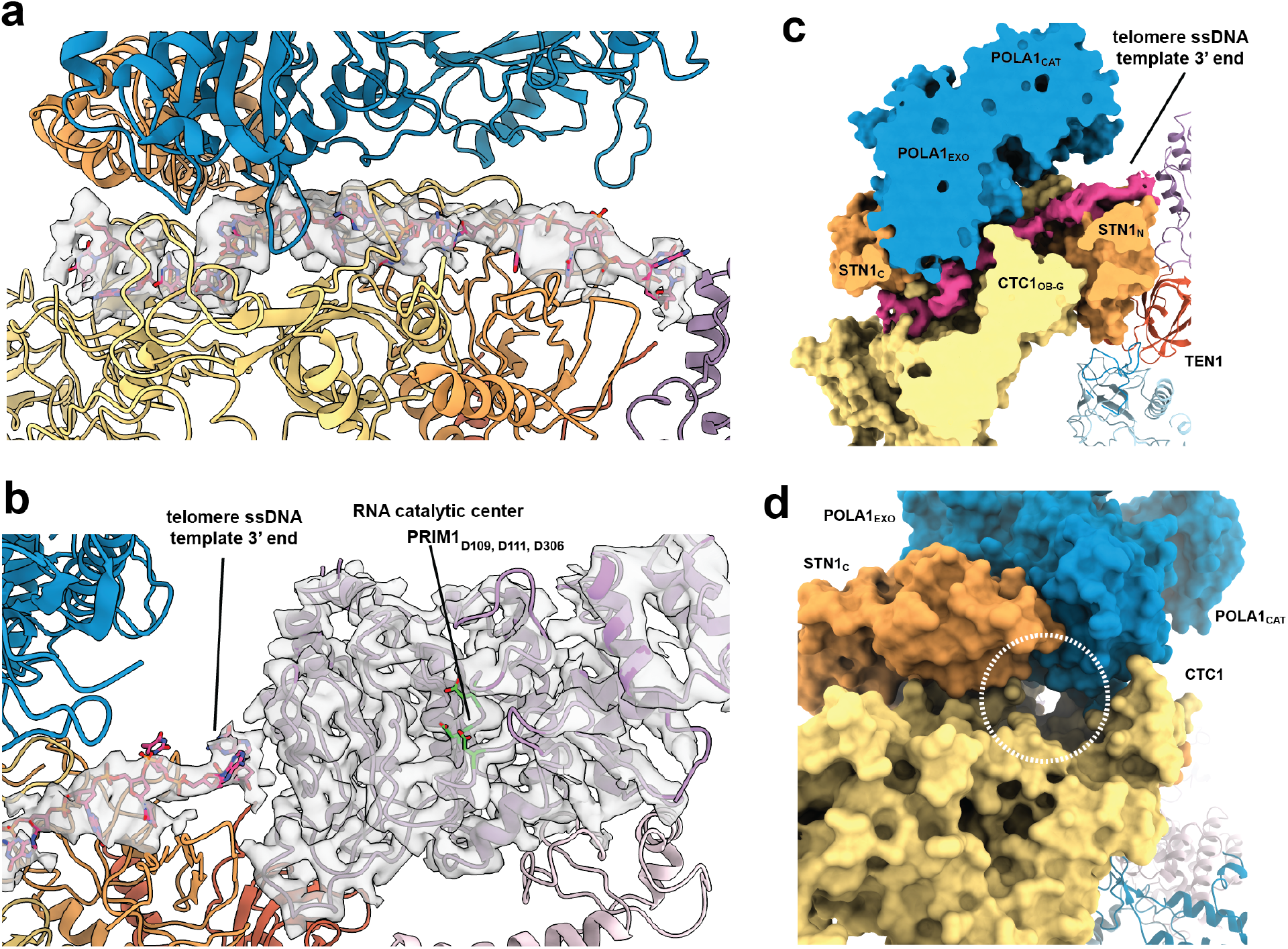
CST poise the ssDNA template for pol-α/primase RNA primer synthesis. **a**, The CTC1 subunit of CST binds the telomeric single-stranded DNA (ssDNA) template. Protein subunits are represented in ribbon cartoons. The modeled ssDNA is shown as pink-colored atomic model while its EM density map is shown in semi-transparent gray volume. STN1_C_ and POLA1_EXO_ sit on top of the CST-ssDNA co-complex. **b**, The 3’ end of the ssDNA is pointing towards the RNA catalytic center of PRIM1. The residues involved in RNA synthesis are colored green. The EM densities of the ssDNA and PRIM1 are shown as semi-transparent gray volumes. **c**, Cross-section view of the molecular tunnel formed by CTC1_OB-G_ and POLA1_EXO_ domains, with the ssDNA threading through. The ssDNA, CTC1, STN1, and POLA1_EXO_ are shown as surface representations. The rest of the subunits are in ribbon cartoons. **d**, The molecular tunnel viewed from the 5’ end of the ssDNA but with the ssDNA model hidden to reveal the tunnel. The entrance of the tunnel is marked by a white dashed circle. The rest of the subunits are in ribbon cartoons. Colors are defined similarly in all panels: CTC1 in yellow, STN1 in orange, TEN1 in dark orange, PRIM1 in purple, PRIM2 in light purple, POLA1 in blue, and POLA2 in light blue.

The catalytically dead exonuclease domain^25,26^ (POLA1_EXO_) of POLA1 sits on top of the ssDNA bound CTC1 (**Fig. 3c**). This is mediated by direct interactions between POLA_EXO_ and the ssDNA and CTC1_OB-G_ (**Supplementary Fig. 8b**). Interestingly, POLA1_EXO_ uses an alpha-helical region to contact the ssDNA template, similar to the yeast POLA1^27^. This region replaces the β-hairpin motif typically found in proofreading B DNA polymerases^25,26^. The STN1_C_ domain further stabilizes the CTC1-POLA1_EXO_ assembly by forming a connecting wedge between the two (**Fig. 2d and 3d**). Thus, CST and POLA1_EXO_ create a tight-fit molecular tunnel for the ssDNA template to thread through (**Fig. 3c & d**). In summary, CST facilitates RNA synthesis initiation using a two-pronged approach - positioning the ssDNA template 3’ end and orienting the PRIM1 RNA catalytic center to face the template 3’ end (**Fig. 2b and 3b**).

### PRIM2_C_ is flexibly tethered and within range to transfer the RNA primer to POLA1_CAT_

The PRIM2 C-terminal domain, PRIM2_C_, facilitates primer initiation by binding the 5’-triphosphate of the primer and coordinating the switch from RNA primer synthesis to DNA elongation^19,21,28,29^. It is connected to the PRIM2_N_ domain by an 18-residue unstructured linker (**Fig. 1a**). The linker length is proposed to confer conformational flexibility for PRIM2_C_ to count the RNA primer length and transfer the “matured” RNA/DNA molecule to POLA1 for DNA elongation^2,29^.

Three-dimensional variability analysis (3DVA)^30^ of our PIC structure reveals the PRIM2_C_ domain exhibits a wide range of motion as compared to the rest of PIC, indicating significant flexibility for this domain (**Fig. 4a, mode 1 and Supplementary Movie 1**). In addition, PRIM2_C_ is positioned in the local vicinity of the 3’ end of the ssDNA template, PRIM1, and POLA1_CAT_ domains (**Fig 4b**). These suggest that PRIM2_C_ is poised to initiate RNA primer synthesis with PRIM1 and within range to coordinate the switch to DNA elongation with POLA1.

**Fig. 4:**
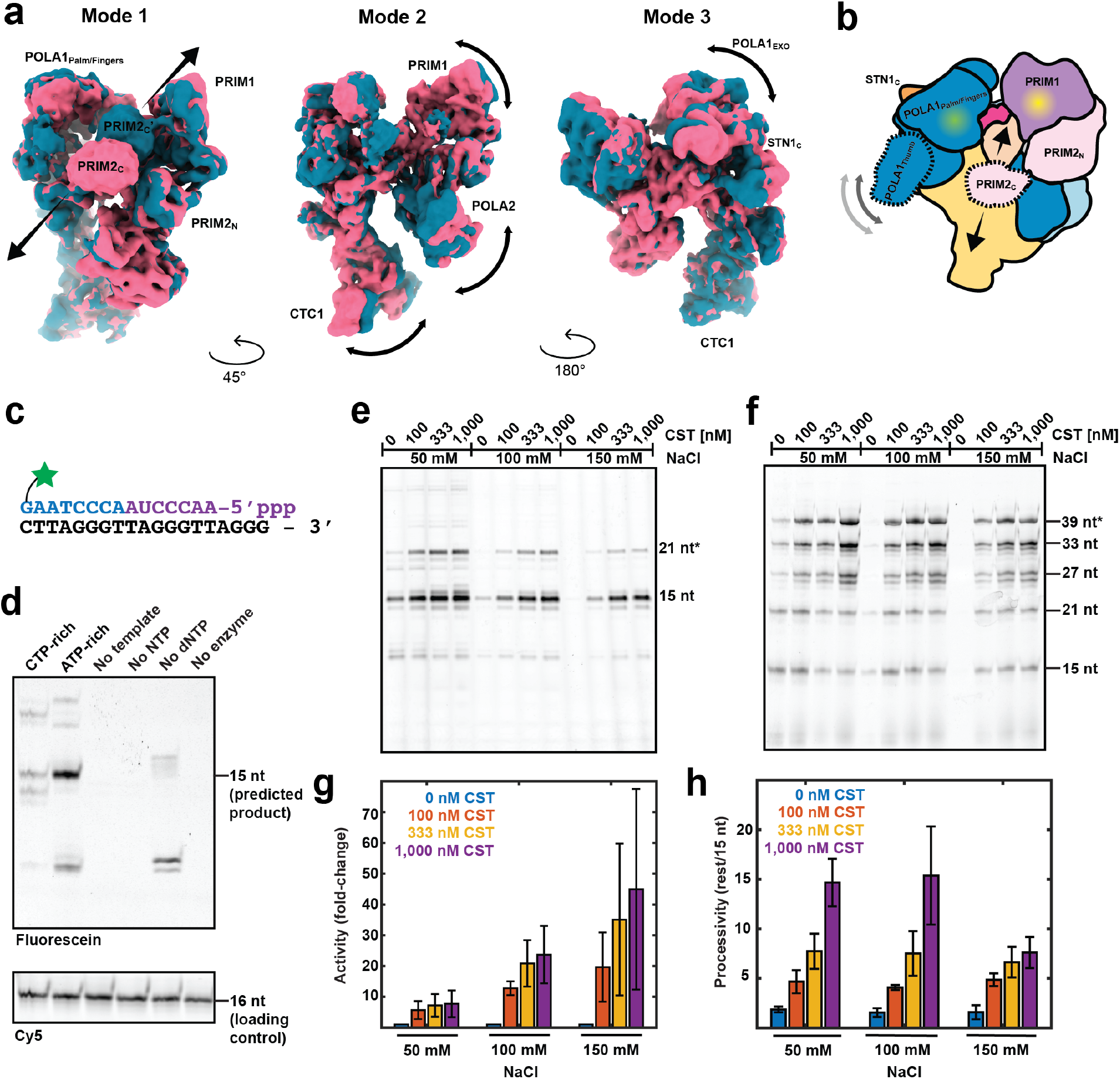
Analysis of the flexibility and enzymatic activity of the PIC complex. **a**, Three-dimension variability analysis (3DVA) of the PIC structure revealed three distinct motions (modes). Each mode is illustrated by two maps that represent the extreme ends of each motion; navy blue and pink maps. Arrows depict the directions of motions. PRIM2_C_ and PRIM2_C_*′* refers to the two termini positions of the domain in the motion. **b**, Cartoon illustration of the motions observed in the 3DVA analysis and flexible (unresolved) domains of pol-α/primase; POLA1_CAT_ thumb subdomains and PRIM2_C_. The RNA and DNA catalytic centers are highlighted with yellow and green spots respectively. The PIC subunits are colored as follows: CTC1 in yellow, STN1 in orange, TEN1 in dark orange, PRIM1 in purple, PRIM2 in light purple, POLA1 in blue, and POLA2 in light blue. **c-h**, fluorescence end-labeled direct assay to study pol-α/primase enzymatic activity. **c**, Predicted RNA-DNA product of pol-α/primase using a C3xTEL ssDNA template (black). RNA and DNA portions of the product are colored in purple and blue respectively. A fluorescein-12-dGTP is used to end-label products. **d**, Reactant depletion experiment to validate that RNA-DNA products are made and detected by the assay. CTP-rich or ATP-rich NTP condition can control initiation start sites, as expected. Unless otherwise indicated, standard reactions used the ATP-rich NTP condition. All reactions used C3xTEL as the DNA template. A Cy5 oligo (16 nt) is used as a loading control and product size reference. **e & g**, CST enhances pol-α/primase total activity. Activity stimulation is tested at different buffer salt conditions and quantified as fold-change over the no-CST sample lanes. **f & h**, CST stimulates pol-α/primase processivity at different buffer salt conditions. Processivity is quantified by total counts above the 15 nt band divided by the counts of the 15 nt band. Asterisks indicate reprimed products.

3DVA also revealed that the PIC exhibits other independent motions in the primase and polymerase lobes (**Fig. 4a, mode 2 & 3**). The POLA1_EXO_ domain is swiveling about its docking position on the ssDNA and CTC1 (**Supplementary Movie 2**). The swiveling motion range is restricted by the flexible peptide linker connecting STN1_C_ and STN1_N_. The analysis also shows the primase lobe is swiveling about the CTC1_OB-G_ domain, independent of the polymerase lobe attached to the rest of CTC1 (**Supplementary Movie 3**).

### CST stimulates both the activity and processivity of pol-α/primase on telomeric templates

Previous studies using poly-dT ssDNA templates showed mammalian CST increases pol-α/primase template affinity and stimulates the enzyme activity and processivity^4,31^, but it is unclear how these findings translate to telomeric ssDNA templates. To study the effects of CST addition to pol-α/primase enzymatic activity on telomeric ssDNA templates, we developed a fluorescence end-labeling coupled primase-polymerase direct assay based on a previous design^32^ (**Fig. 4c & d**). In an ATP-rich NTP condition (as in mammalian cells^33^), pol-α/primase initiates the rate-limiting step of dinucleotide synthesis^34^ at every double thymidine position of a telomeric DNA template. This yields ladder-like products spaced six nucleotides (nt) apart due to the repetitive hexameric telomeric sequence^32^.

Using this assay, we found pol-α/primase made a major 15 nt RNA-DNA product when a 3-repeat telomeric ssDNA template is used (C3xTEL, 19 nt, the same template that we used in our cryo-EM experiments) (**Fig. 4d & e**). The addition of CST stimulates pol-α/primase activity multiple-fold (**Fig. 4e & g**). This result corroborates past studies using poly-dT templates^4,31^. With a longer telomeric ssDNA template that produces ladder-like products, we could indirectly quantify pol-α/primase processivity changes (end-product processivity). Using a 6-repeat telomeric ssDNA template (C6xTEL, 37 nt), CST addition stimulates pol-α/primase processivity by ∼ 10-fold (**Fig. 4f & h**). All the above demonstrate that our cryo-EM structure represents an active preinitiation co-complex that can progress to RNA primer synthesis and primer DNA elongation.

### Implications for pol-α/primase lagging-strand DNA synthesis at telomeres and genome-wide

Biochemical studies revealed that CST enhances pol-α/primase affinity to DNA templates and stimulates the enzyme processivity^31,35^, but it was unclear how these functions are performed. Our PIC structure provides a structural basis to explain how CST, as an ssDNA-binding accessory factor, can elevate pol-α/primase affinity to ssDNA templates and stimulate enzyme activity. First, CST binds and stretches out the ssDNA template. The CST-ssDNA complex then provides a scaffold for pol-α/primase assembly, allowing it to change from a compact apo-state to a bipartite form with spatially separated primase and polymerase domains (**Fig. 2a**). Because CST binds the ssDNA template, its ssDNA-binding preference would dictate the genome location of pol-α/primase recruitment.

Our PIC structure shows that pol-α/primase assembly with CST-ssDNA leads to a conformation that geometrically favors primase-template binding for *de novo* RNA primer synthesis. CST does this via a two-pronged approach; by presenting the 3’ end region of the ssDNA template to PRIM1 and orientating the RNA catalytic face of PRIM1 towards the template. Although our PIC structure shows the molecular 3’ end of the template pointing towards the PRIM1 RNA catalytic center, we expect our model to be compatible with a longer template where the co-complex acts on an internal ssDNA region with the strand polarity still preserved. This is supported by our observations of similar-looking PIC negative-stain EM 2D class averages when longer telomeric ssDNA templates were used (**Supplementary Fig. 4**).

The PRIM2_C_ domain is critical for the pol-α/primase RNA-to-DNA switch mechanism. It binds the RNA primer^21,29^ and hands it over to POLA1 for DNA elongation^34,36^. These processes hinge on the 18-residue flexible linker^37^ (**Fig. 1a**) connecting PRIM2_C_ to PRIM2_N_. However, it was previously unclear how these findings coalesce into a structural model. This work shows that the PRIM2_C_ domain is indeed highly flexible. More importantly, the PIC architectural organization positions the domain within reach of either the RNA or DNA catalytic center (**Fig. 4a - mode 1 & 4b and Supplementary movie 1**). Thus, our PIC structure provides a model that rationalizes previous biochemical findings and shows that the PRIM2_C_ domain is poised to facilitate RNA primer initiation, elongation, and primer hand over to POLA1. In other words, our PIC structure reveals a single platform architecture that pol-α/primase and single-stranded DNA-binding accessory factors could form to mediate the multi-steps *de novo* primer synthesis process. Given the structural and biochemical similarity between CST and RPA^5,7^, it will be interesting to investigate if RPA and pol-α/primase form a similar PIC structure at canonical replication forks.

## Methods

### Cloning

Human *POLA1* (NP_058633.2) and *POLA1*_Δ1-337_ was cloned into the pOET1 transfer vector (Mirus Bio). Human *PRIM1* (NP_000937.1), *PRIM2* (NP_000938.2) and *POLA2* (NP_002680.2) cDNAs were individually cloned into the pFastBac1 expression vector (Invitrogen). The POLA1 and POLA1_Δ1-337_ expression vectors have N-terminal twin-strep tag while that of POLA2, PRIM1 and PRIM2 constructs have N-terminal hexa-histidine (6xHis) tags. The human CST subunits cDNA, *CTC1* (AAI11784), *STN1* (NP_079204) and *TEN1* (NP_001106795) were cloned into a single pBAC4x-1 transfer vector (Novagen). CTC1 open reading frame has a N-terminal 3xFLAG while both STN1 and TEN1 have 6xHis tags.

### Insect cells culture and baculovirus generation

Recombinant baculovirus encoding POLA1 (or its truncation mutant) were produced using the flashBAC ultra system (Mirus Bio). The baculoviruses encoding PRIM1, PRIM2 and POLA2 were made using the Bac-to-Bac system (Invitrogen). The viruses were amplified to a tier of > 1.0 × 10^8^ pfu/mL before using them for insect cells infection. The virus titers were measured by flow cytometry assay (Expression Systems).

### Expression and purification of CST and pol-α/primase

One or two liters of *Trichoplusia* (Tni) cells (Expression System) were infected with recombinant baculoviruses at a cell density of 1.5-2.0 × 10^6^ cells per mL. A single baculovirus stock was used for CST expression while four baculoviruses (POLA1, POLA2, PRIM1 and PRIM2) were used to co-express DNA polymerase-alpha/primase complex (pol-α/primase). Either infection was done at a multiplicity of infection of one. The infected Tni cells were further incubated in an orbital shaker (27 °C, 130 r.p.m.) for 66-68 h before harvesting.

CST protein complex was purified based on an established protocol^7^. The infected Tni cells were harvested by centrifugation at 1,500 x g for 30 min at 4 °C. The cell pellets were resuspended in lysis buffer (25 mM HEPES-NaOH pH 7.5, 300 mM NaCl, 15 mM imidazole, 1 mM dithiothreitol (DTT), 1x Xpert Protease Inhibitor Cocktail Solution (GenDEPOT)) at 50 mL lysis buffer per liter of cells and subjected to ultrasonication for cellular lysis. The cell lysate was clarified by high-speed centrifugation and the clarified lysate incubated with Ni-NTA agarose resin (Qiagen) for 2 h at 4 °C using a rotator. The resin was washed three times with wash buffer (same as lysis buffer but without protease inhibitors) and eluted with Ni-NTA elution buffer (25 mM HEPES-NaOH pH 7.5, 300 mM NaCl, 300 mM imidazole, 1 mM DTT). The elute was then incubated with anti-FLAG resin (GenScript) overnight at 4 °C. The resin was washed three times with wash buffer and the CST complex eluted with FLAG elution buffer (25 mM HEPES-NaOH pH 7.5, 300 mM NaCl, 1 mM DTT, 0.4 mg/mL 3xFLAG peptide (APExBIO)).

Pol-α/primase purification is same as the CST purification except the buffers contain 150 mM NaCl instead of 300 mM NaCl and that a twin-strep tag pull-down was performed instead of a FLAG tag pull-down. The Ni-NTA elute was incubated with Strep-Tactin XT 4Flow-resin (IBA LifeScience) overnight at 4 °C. The resin was washed three times with wash buffer (25 mM HEPES-NaOH pH 7.5, 150 mM NaCl, 1 mM DTT) before eluting the Pol-α/primase complex with 1x BXT elution buffer (IBA LifeScience). The purity and integrity of both protein complexes were checked with SDS-PAGE analysis before they are concentrated to ≥ 5.0 mg per mL. The complexes were used in experiments or snap-frozen in small aliquots for long-term storage in −80 °C.

### Co-complex formation

The purified CST and pol-α/primase were mixed at a 2:1 molar-ratio and incubated at 4 °C overnight. Strep-Tactin XT 4Flow-resin was added to the mixture and incubated for 1 hour at 4 °C. The resin was washed four times with wash buffer (25 mM HEPES-NaOH pH 7.5, 150 mM NaCl, 1 mM DTT) before the co-complex was eluted with 1x BXT elution buffer (0.5 h incubation). The integrity and purity of the co-complex were confirmed with SDS-PAGE before experimental use or snap-frozen for long-term storage in −80 °C.

### Pull-down assay

The protein bait and prey samples were incubated for 1 h at 4 °C before Strep-Tactin XT 4Flow-resin was added to capture the twin-strep-tagged bait proteins. The resin was washed three times with wash buffer before the captured samples eluted with 1x BXT elution buffer (incubated for 0.5 hour). The pulled down protein samples were analyzed with Coomassie-stained SDS-PAGE.

### Agarose gel-based binding assay

Radioactive ^32^P 5′-labeled oligonucleotides were used in the binding assays. The estimated specific activity was at least 200,000 c.p.m. per pmol, and 500 c.p.m was used for each reaction. For binding, purified protein complexes and DNA were mixed and incubated as a 10 µL reaction volume in binding buffer (50 mM HEPES-NaOH pH 7.5, 150 mM NaCl, 5 mM MgCl_2_, 10 % glycerol, 0.1 mg/mL BSA, 2 mM TCEP) for 1 h at 25 °C. The samples were then loaded into a pre-chilled 1x TBE 0.7 % agarose gel and the gel ran at 7 V/cm for 1.5 h in a cold room (4 °C). The gels were vacuum dried at 80 °C for 1.5 h before exposing to a storage phosphor screen overnight. The screen was then imaged with a Typhoon FLA 9000 scanner (GE Lifesciences, USA). Binding analysis was done using Fiji (National Institutes of Health) and curve fitting performed with GraphPad Prism (GraphPad Software).

### Negative-stain electron microscopy sample preparation, data collection and image analysis

Samples were immediately diluted to 75-100 nM before deposition to glow discharged electron microscopy (EM) grids (CF400-Cu-UL, EMS). The grids are then subjected to 2 % (w/v) uranyl formate (Structure Probe) staining using an established protocol^38^. The negative-stain EM sample grids were imaged on a 120 kV transmission electron microscope (Talos L120C, Thermo Fisher Scientific). Image datasets were collected with SerialEM^39^ using low dose mode. Two-dimensional (2D) classification analysis was performed using CryoSparc2^40^.

### Cryogenic-electron microscopy sample preparation and data collection

Cryo-EM sample grids for the co-complex were prepared as described for CST^7^. Briefly, 1.2/1.3 µm 300 mesh holey grids (Quantifoil) were glow discharged for 20 s at 15 mA (PELCO easiGlow) before use. Proteins samples were then added to the freshly glow-discharged grids and vitrified using Vitrobot (Thermo Fisher Scientific) with 4.5 s blot time at 4 °C and 95 % relative humidity.

Suitable cryo-EM sample grids for data collection were first screened on an Arctica 200 kV transmission electron microscope (Thermo Fisher Scientific) that is equipped with a Bioquantum K3 energy filter-detector (Gatan). A dataset (7,470 movies) of CST-pol-α/primase-C3xTEL sample grid was collected using SerialEM on the Arctica with a calibrated pixel size of 1.064 Å/pixel, a defocus range of −0.5 to −2.5 µm, a total dose of 48.5 e^-^/Å^2^ (40 frames), and the K3 detector (20 eV energy slit) in counting mode.

### Cryogenic-electron microscopy single-particle analysis

All movie frames (7,470 movies) were motion corrected (5 × 5 patch)^41^ and their CTF^42^ estimated in Relion. A total of 1,156,310 particles were picked with Relion Laplacian-of-Gaussian (LoG) auto-picker. The particles were extracted with a 4x binning (4.256 Å/pixel) and exported to CryoSparc2^40^ for two rounds of four-class hetero-refinement (initial models populated from ab initio modeling step) and two rounds of two-dimensional classification (100 classes). The resulting 245,828 particles were subjected to a four-class ab initio modeling step to further isolate the co-complex structure. Similar-looking classes were combined (236,905 particles) and progressed to non-uniform refinement^43^. The particles were re-extracted at its original pixel size (1.064 Å/pixel) and with re-centering done using the poses from the prior 3D refinement. The newly extracted particles were used for 3D auto-refinement in Relion which resulted in an initial 4.78 Å map. Using the new refined poses, one round of particle polishing and two rounds of CTF refinements were performed in Relion^41,44^. A second 3D auto-refinement step was performed yielding a 4.43 Å map. Additional structural heterogeneity was further sieved by subjecting the particles to a six-class 3D classification step in Relion with no alignments performed. The classification isolated a single class (∼67%) consisting of 159,979 particles that was exported to CryoSparc2 for non-uniform refinement, yielding a 4.00 Å map. An additional round of local CTF refinement in CryoSparc2 and non-uniform refinement resulted in the reported map of 3.86 Å global resolution. The conversion of files from that of CryoSparc2 to Relion was performed using UCSF PyEM scripts (https://github.com/asarnow/pyem).

### Model building, refinement, and validation

All seven protein subunits of the preinitiation complex (PIC) were built using computed models from the AlphaFold Protein Structure Database (https://alphafold.ebi.ac.uk/). The models were first manually moved into their respective positions in the 3.86 Å cryo-EM map and then flexibly fitted using the Jiggle fit function in Coot^45^. The single-stranded DNA molecule was built starting from the four nucleotide model shown in the previous CST structure^7^. The assembled PIC model was visually checked in Coot before subjecting to real-space refinement in PHENIX^46^. Model to map validations were performed using PHENIX cryo-EM comprehensive validation program

### Enzyme direct assays

A standard reaction (20 μL) consists of NTPs, dNTPs, 5 μM DNA template, and 500 nM pol-α/primase in direct assay buffer (of 50mM of HEPES-NaOH pH7.5, 100 mM NaCl, 5 mM MgCl_2_, and 2 mM DTT). Addition of CST and variation in NaCl concentrations are as indicated in figure legends or annotations. Otherwise stated, the NTPs used was ATP-rich; 2 mM of ATP, 50 μM of UTP and CTP each. To visualize the end products, the DNA template has a 5’ C and the dNTPs included 0.5 μM of fluorescein-12-dGTP (AAT Bioquest) in addition to 50 μM of dATP, dTTP and dCTP. DNA templates of various telomeric repeat C(TTAGGG)_n_ were ordered from vendor (IDT). The enzymatic reactions were initiated by last adding the DNA template. The reactions were incubated at 37 °C for 1 h before they were quenched by DNA precipitation. A Cy5-labeled oligo was added as a loading control in this step. After DNA precipitation, each sample was dissolved in 20 μL 1X formamide loading dye and 10 μL of it loaded into each well of a 23 % 1x TBE 7 M Urea PAGE gel. Electrophoresis was performed at a constant 25 W until the bromophenol blue dye reached a third of the way from the gel bottom. The gels were imaged on a Typhoon FLA9000 gel imager using dual fluorescence channels (Fluorescein and Cy5, each at 450 V PMT). Enzyme activity and processivity values were quantified using Fiji, as described in the respective figure legends.

## Acknowledgments

We like to thank Dr. Tahir Tahirov and Dr. Andrey Baranovskiy from UNMC for their initial discussion and help in setting up our experiments. We thank Dr. Tom Cech and Dr. Arthur Zaug from CU Boulder for help discussion and suggestions. We also like to thank our colleagues, Dr. Samuel Butcher and Dr. Judith Kimble for their input, discussion, and critical reading of the manuscript. We also like to thank the members of the Lim lab for their helpful discussions and suggestions. We are grateful for the use of facilities and instrumentation at the Cryo-EM Research Center in the Department of Biochemistry at the University of Wisconsin-Madison. Support for this research was provided to C.L. by the National Institutes of Health (NIH), National Institute of General Medical Sciences (NIGMS) (R00GM131023), and the University of Wisconsin–Madison, Office of the Vice-Chancellor for Research and Graduate Education with funding from the Wisconsin Alumni Research Foundation and the Department of Biochemistry.

## Author contributions

Q.H. and C.L. designed the expression constructs for recombinant proteins production, with support from B.L., Q.H. and X.L. performed insect cell culture and baculovirus production for insect cell infection. Q.H. expressed and purified the recombinant human protein complexes. Q.H., B.C, and C.L. conducted the negative-stain EM data collection and sample screening. Q.H. and C.L. prepare the cryo-EM samples for data collection. C.L. processed and analyzed the cryo-EM datasets with support from Q.H. and B.C.. C.L. and Q.H. performed model building and refinement. C.L. and X.L. performed the gel-based enzyme assay, and C.L., X.L., and Q.H. analyzed the data. C.L. and B.C. performed the EMSA assays, and X.L., B.C., and Q.H. analyzed the data. C.L. directed the project and designed the experiments with Q.H., X.L., and B.C.. C.L. wrote the manuscript with inputs from all authors.

---

**Supplementary Fig. 1:**
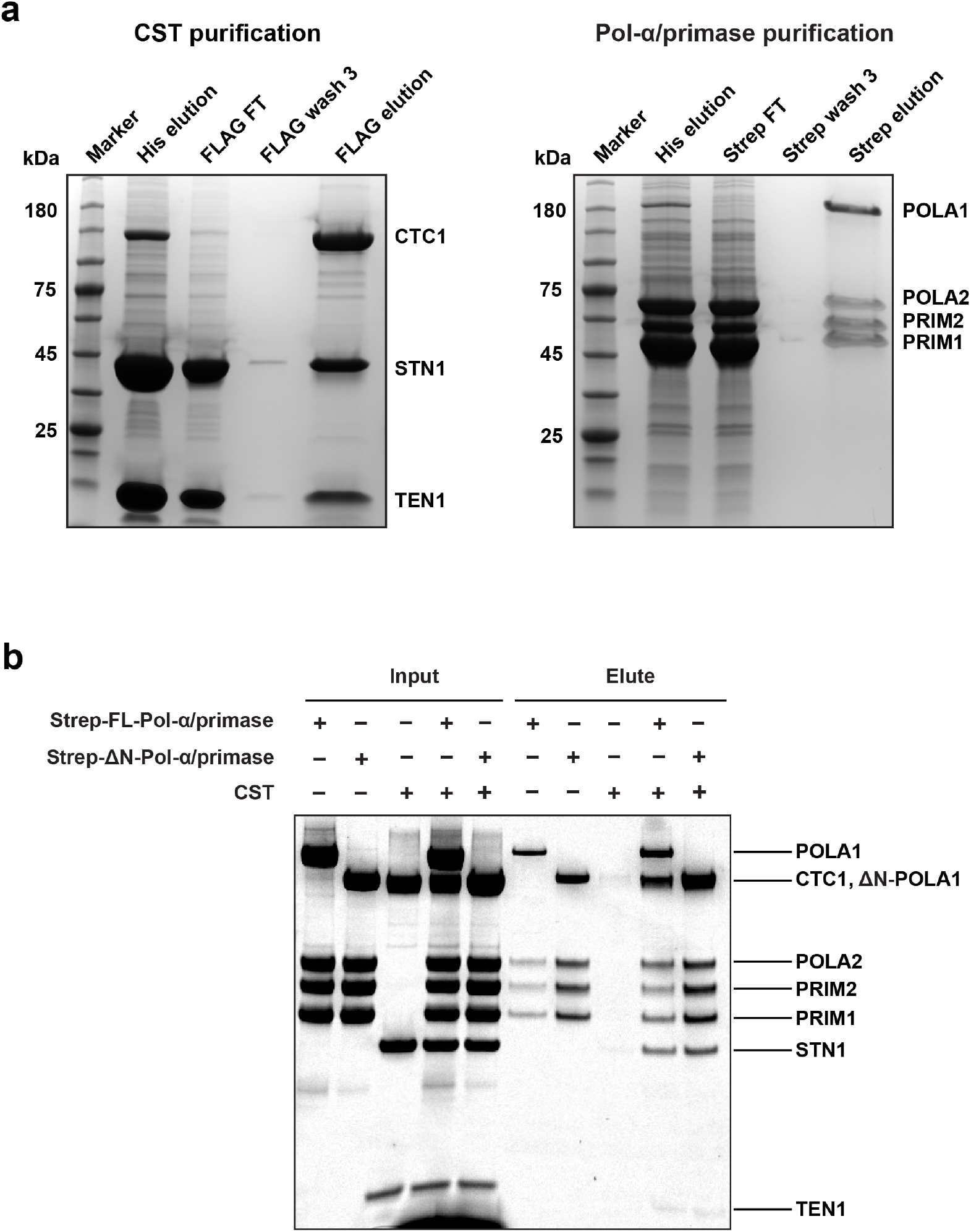
CST and pol-α/primase purification and pull-down assay. **a**, Tandem affinity tags purification of CST and pol-α/primase complexes from baculovirus-infected insect cells. CST heterotrimeric complex was purified by first using Ni-NTA beads to pull down 6xHis-STN1 and 6xHis-TEN1 and then using anti-FLAG beads to pull-down 3xFLAG-CTC1. Pol-α/primase 4-subunit complex was purified by first using Ni-NTA beads to pull down 6xHis-PRIM1, 6xHis-PRIM2, and 6xHis-POLA2 and then using Strep beads to pull-down Strep-POLA1. FT: flowthrough. Trident (GeneTex) high range prestained protein marker was used in both coomassie-stained denaturing protein gels. **b**, Pull-down assay to characterize direct protein-protein interactions between CST and full-length or truncated (ΔPOLA1_1-337_) pol-α/primase (ΔN-pol-α/primase). Strep beads were used with pol-α/primase as bait and CST as prey. All input lanes are loaded as 10% of total sample.

**Supplementary Fig. 2:**
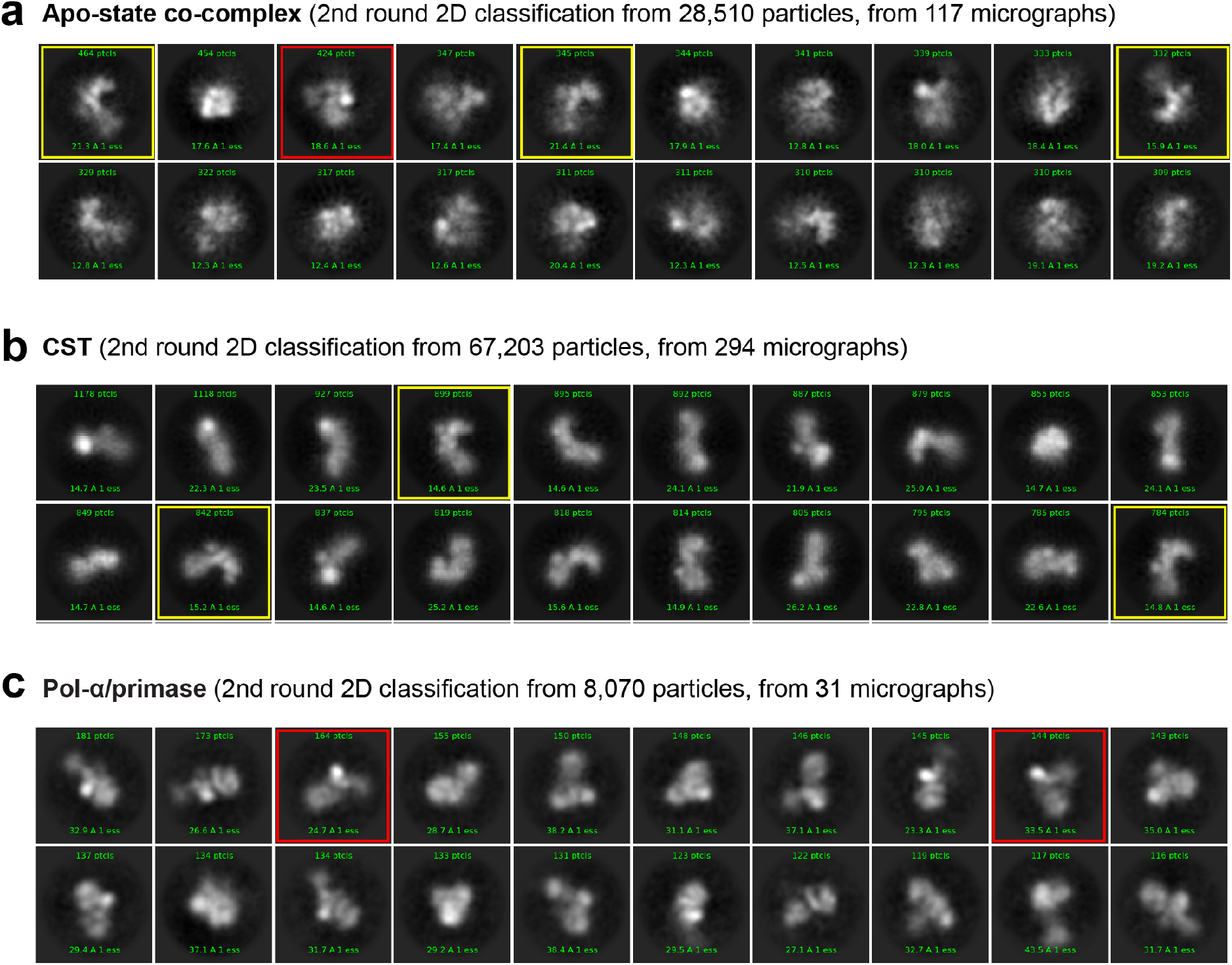
Negative-stain electron microscopy 2D classification analysis of CST and pol-α/primase apo-state co-complex. **a-c**, Top twenty (by distribution) 2D class averages of each apo-state CST-pol-α/primase, CST and pol-α/primase. **a**, No new and distinct 2D class averages are seen in the apo-state CST-pol-α/primase analysis. Instead, class averages similar to that of CST alone (**b**) or pol-α/primase (**c**) are seen. These class averages are indicated with yellow boxes for CST and red boxes for pol-α/primase.

**Supplementary Fig. 3:**
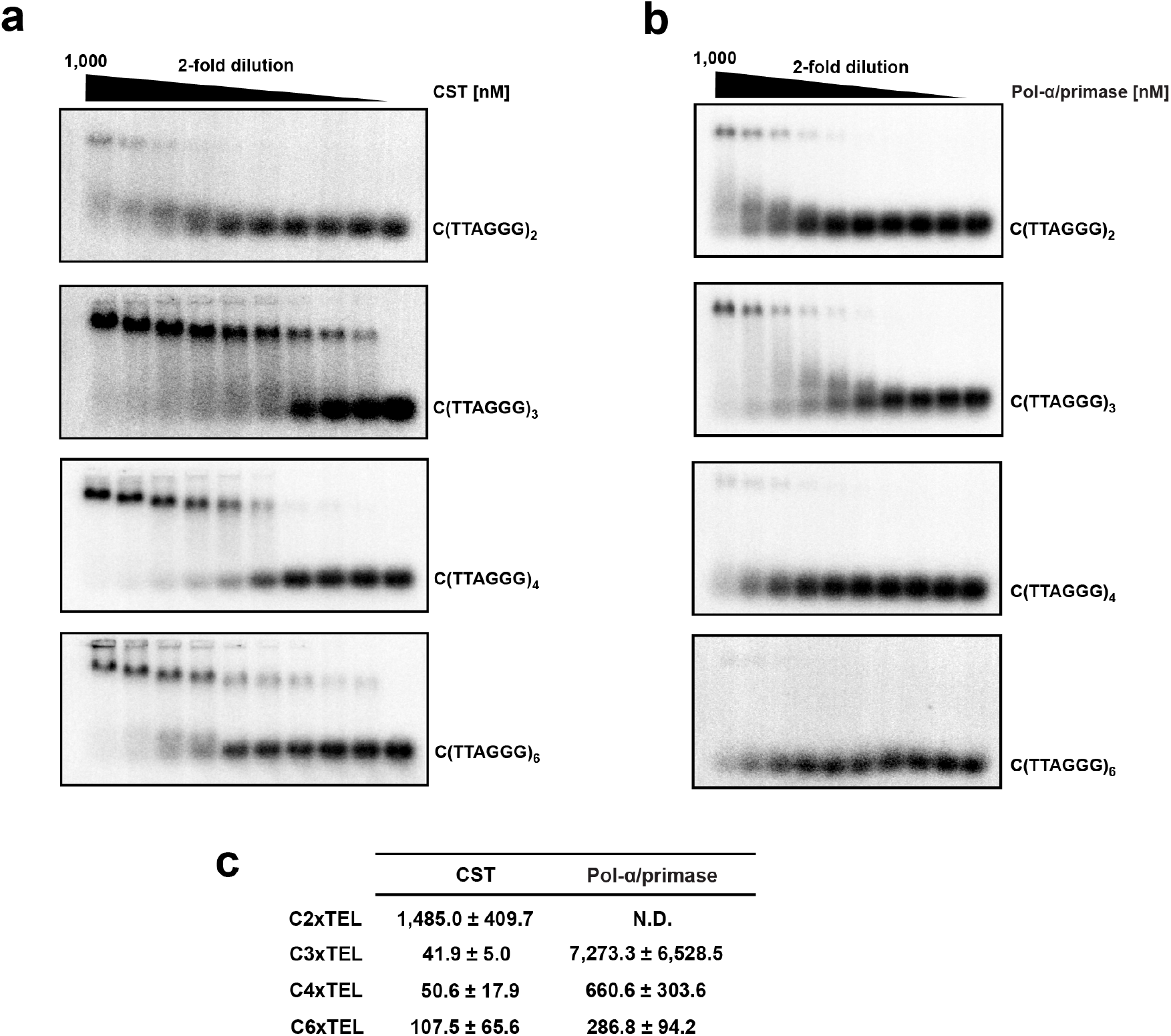
Electrophoretic mobility shift assay to study telomeric single-stranded DNA binding affinity of CST and pol-α/primase. **a**. Representative agarose-EMSA showing CST binds to various length of telomeric single-stranded DNA, from 2 to 6 repeat of TTAGGG. **b**. Same as **a**, but with pol-α/primase. **c**. The binding affinity (*K*_d,app_) was determined by curve fitting (single binding site Hill equation) through quantification of band intensities from the gels. The reported values are averages while the errors represent standard deviations (*n* = 3 independent experiments). N.D. means binding was too weak for curve fitting.

**Supplementary Fig. 4:**
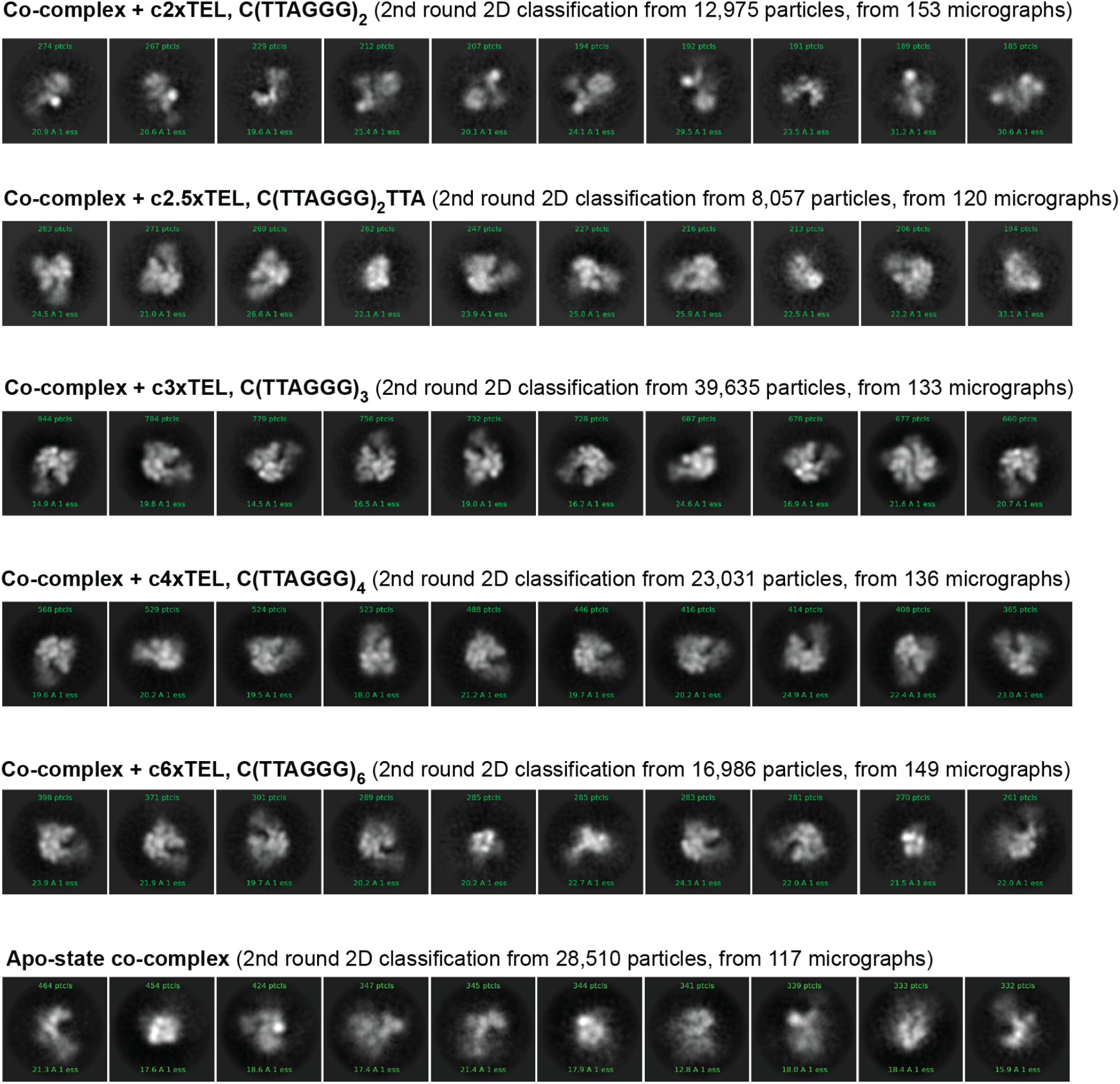
Negative-stain electron microscopy 2D classification analysis of CST and pol-α/primase co-complex bound to telomeric single-stranded DNA. Top ten (by distribution) class averages of CST-pol-α/primase preincubated with various length of telomeric single-stranded DNA, from 2 to 6 repeat of TTAGGG. Every telomeric ssDNA has a 5’ C for consistency with our enzymatic direct assays. The apo-state co-complex analysis served as a no-DNA control. The c2xTEL ssDNA sample has the same outlook as that without ssDNA (apo-state). We see similar 2D class averages when c2.5xTEL to c6xTEL ssDNA were added to the co-complex.

**Supplementary Fig. 5:**
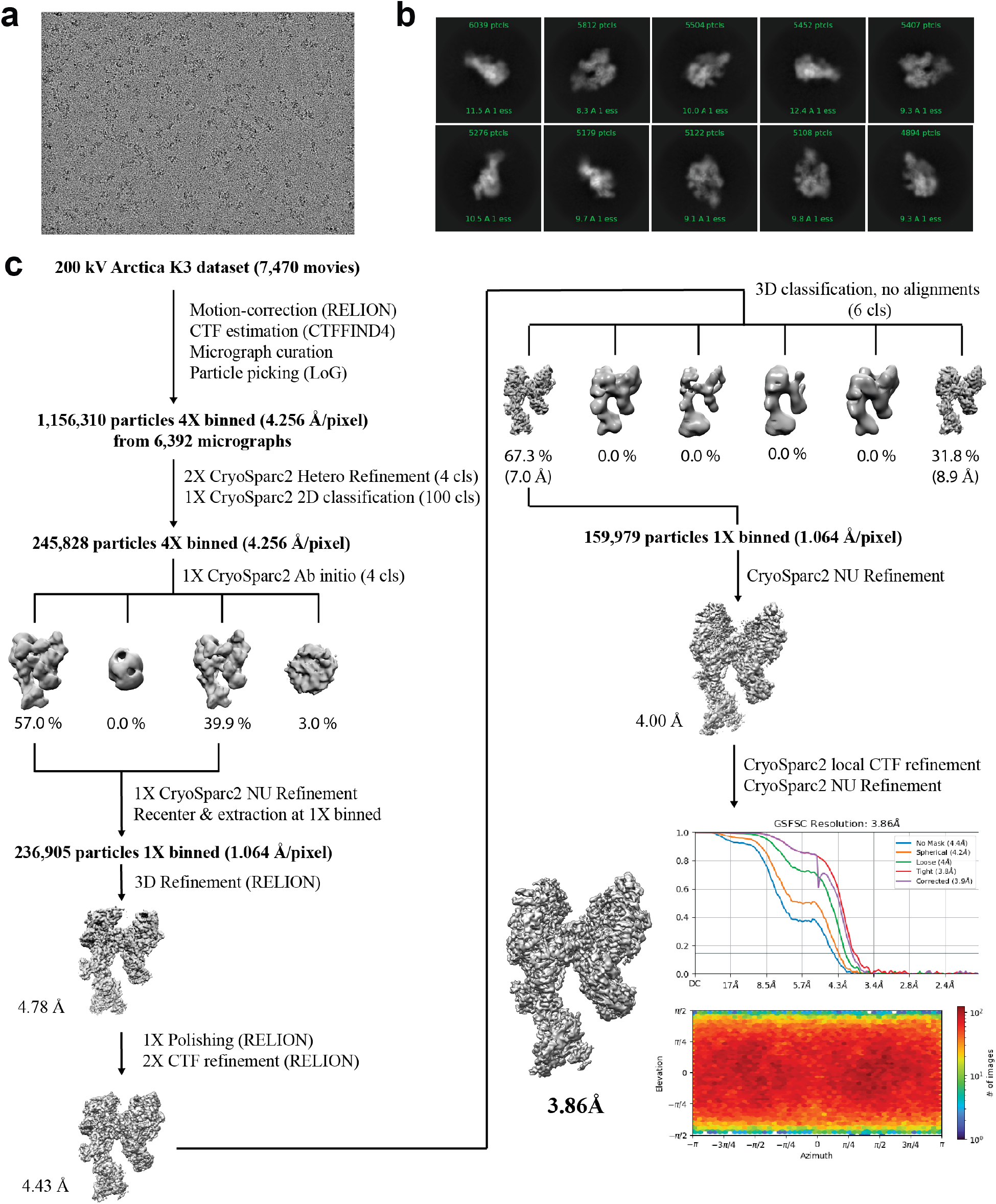
Cryo-electron microscopy data processing pipeline for PIC. **a**, Representative motion-corrected micrograph. **b**, Top ten 2D class averages of PIC. **c**, processing pipeline of the PIC dataset. Class percentages were rounded down to the first decimal place.

**Supplementary Fig. 6:**
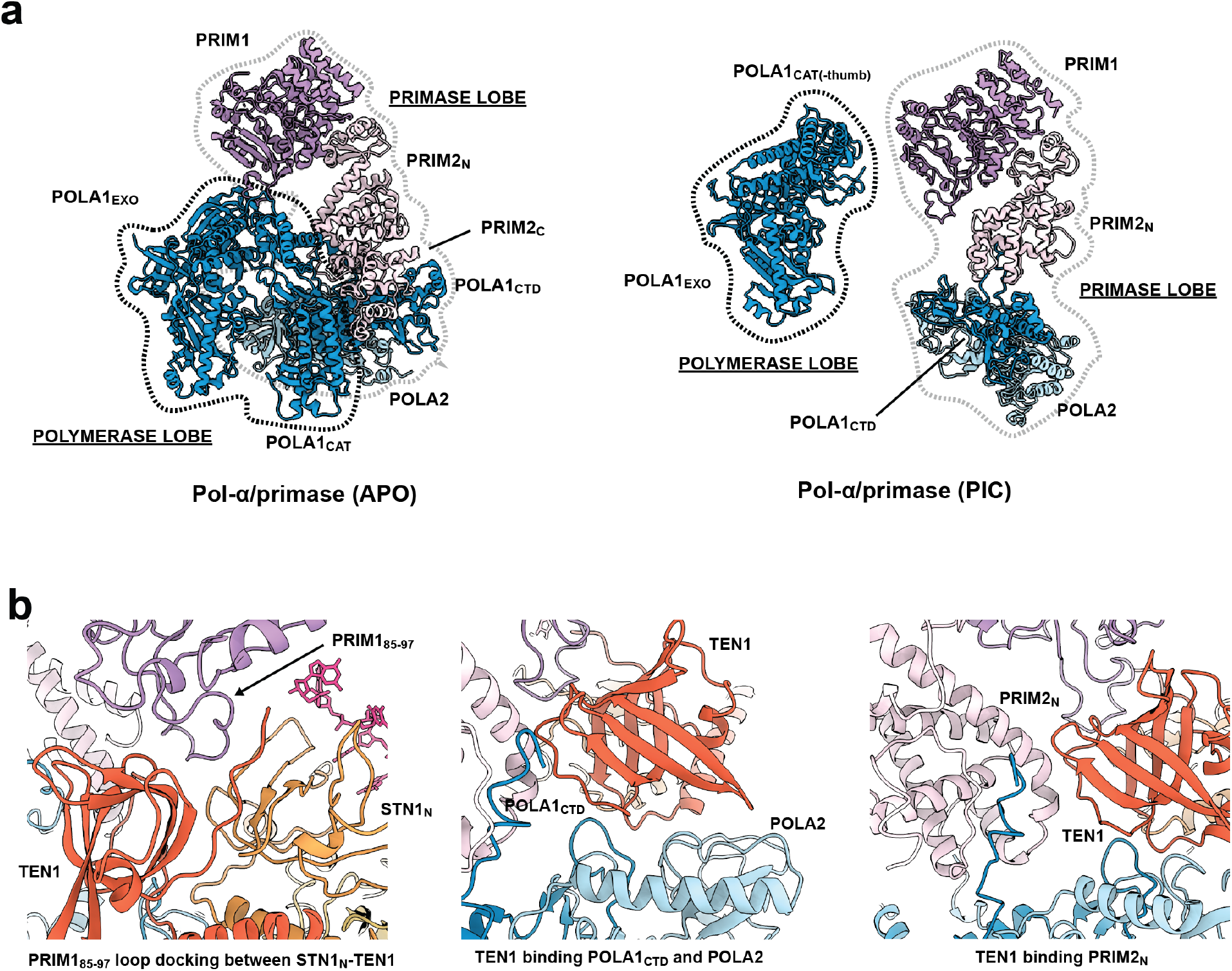
Conformation change in pol-α/primase architecture from apo- to preinitiation-state. **a**, Pol-α/primase architecture changes from a compact shape to a segregated form upon forming the preinitiation complex (PIC) with CST-ssDNA. The APO and PIC structures are depicted as ribbon cartoons. CST and ssDNA are hidden in the PIC structure to better demonstrate the segregated architecture of the PIC structure – separated into polymerase and primase lobes. The lobe boundaries are loosely defined by the dashed lines – black for polymerase lobe and grey for primase lobe. POLA1_CAT_ thumb subdomain and PRIM2_C_ domain are not illustrated in the PIC structure because of their flexibility (see main text). **b**, Zoomed-in representations of the primase lobe interaction with STN1_N_ and TEN1 subunits of CST upon PIC assembly. Colors are defined similarly in all panels: CTC1 in yellow, STN1 in orange, TEN1 in dark orange, PRIM1 in purple, PRIM2 in light purple, POLA1 in blue, and POLA2 in light blue. The APO model is adapted from PDB 5EXR.

**Supplementary Fig. 7:**
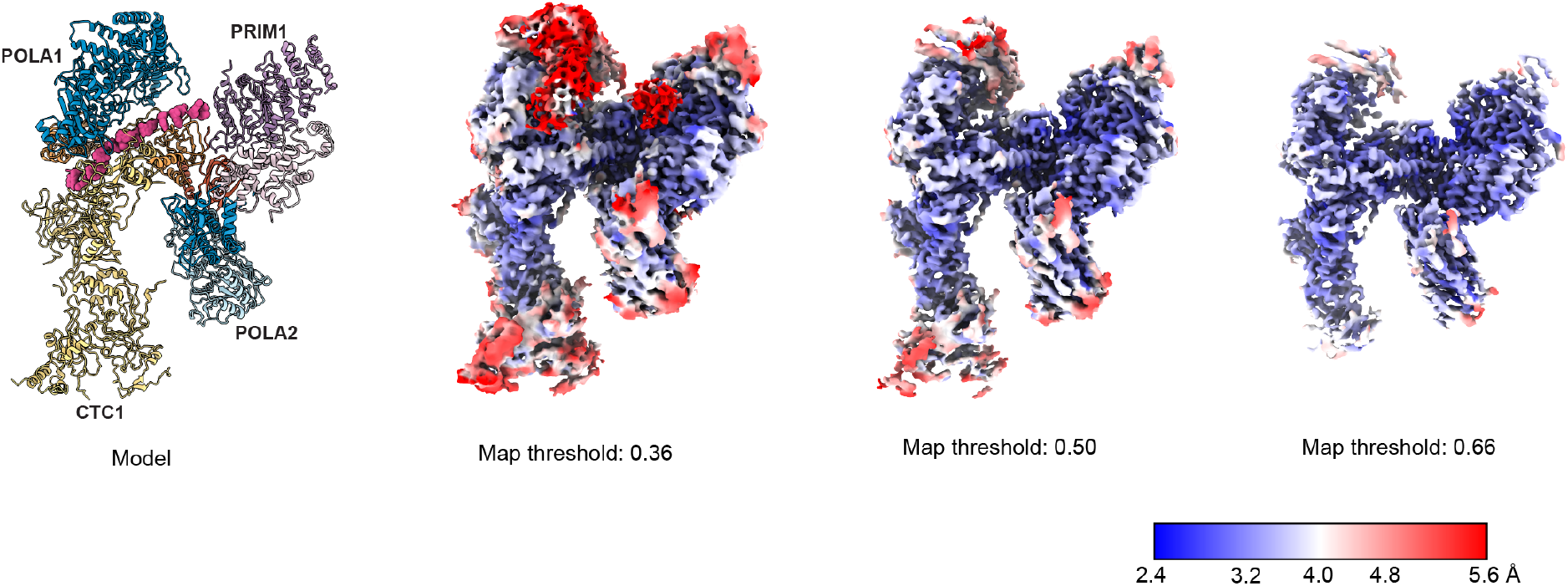
Local resolution map of PIC cryo-EM structure at different map thresholds. Local resolution map of the cryo-EM of the PIC structure. The model is provided as a guide to refer to protein subunit regions on the cryo-EM map. The cryo-EM map is illustrated at various volume thresholds. Color key to local resolution values is shown at bottom right. The model is colored as following: CTC1 in yellow, STN1 in orange, TEN1 in dark orange, PRIM1 in purple, PRIM2 in light purple, POLA1 in blue, and POLA2 in light blue.

**Supplementary Fig. 8:**
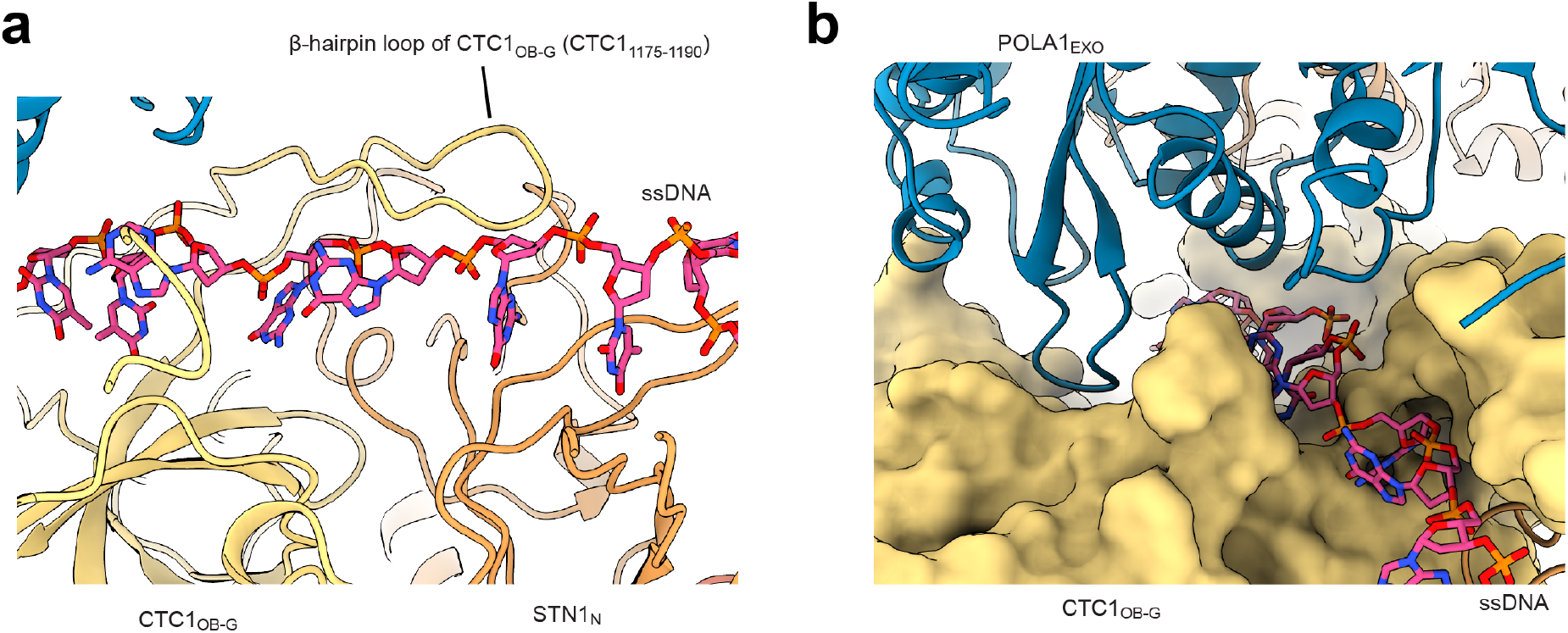
Interaction sites of POLA1 and CST with telomeric ssDNA. **a**, The β-hairpin loop of CTC1_OB-G_ (CTC1_1175-1190_) and STN1_N_ support the 3’ end portion of the ssDNA template. **b**, POLA1_EXO_ domain contacts both CTC1_OB-G_ and the ssDNA template.

**Supplementary Table 1:** Cryo-EM model refinement and map statistics Cryo-EM data collection, refinement and validation statistics.

| | CST-pol-α/primase-C3xTEL |
| --- | --- |
| **Data collection and processing** |  |
| Magnification | 79,000x |
| Voltage (kV) | 200 |
| Electron exposure (e-/Å<sup>2</sup>) | 48.5 |
| Defocus range (μm) | -0.5 to -2.5 |
| Pixel size (Å) | 1.064 |
| Symmetry imposed | C1 |
| Initial particle images (no.) | 1,156,310 |
| Final particle images (no.) | 159,979 |
| Map resolution (Å) | 3.86 |
| FSC threshold | 0.143 |
| Map resolution range (Å) | 2.349 to 10.452 |
| **Refinement** |  |
| Initial model used (PDB code) | AlphaFold |
| Model resolution (Å) | 3.8 |
| FSC threshold | 0.143 |
| Model resolution range (Å) | 3.7 to 4.1 |
| Map sharpening *B* factor (Å<sup>2</sup>) | -171 |
| Model composition |  |
| Non-hydrogen atoms | 28749 |
| Protein residues | 3552 |
| Ligands (DNA) | 15 |
| *B* factors (Å<sup>2</sup>) |  |
| Protein | -218.12 |
| Ligand (DNA) | -177.16 |
| R.m.s. deviations |  |
| Bond lengths (Å) | 0.003 |
| Bond angles (°) | 0.692 |
| Validation |  |
| MolProbity score | 2.31 |
| Clashscore | 14.57 |
| Poor rotamers (%) | 0 |
| Ramachandran plot |  |
| Favored (%) | 85.93 |
| Allowed (%) | 13.33 |
| Disallowed (%) | 0.74 |

